# Entering new circles: Expansion of the LC8/DYNLL1 interactome in the ciliary-centrosomal network through system-driven motif evolution

**DOI:** 10.1101/2023.09.15.557927

**Authors:** Mátyás Pajkos, Tamás Szaniszló, Máté Fülöp, Zsuzsanna Dosztányi

## Abstract

LC8 is a eukaryotic hub protein that presents a highly conserved interface recognized by short linear motifs (SLiMs) in partner proteins. LC8 was originally associated with the dynein motor complex and was also suggested to promote dimerization of partner proteins. However, the growing list of validated partners with diverse functions suggests that LC8 plays a more general cellular role which is still not fully understood. In this work, we combined computational and experimental approaches in order to gain insights into the LC8 interaction network at the system level. Our machine-learning based pipeline together with functional enrichment analysis revealed that LC8 plays a central role in the ciliary-centrosomal system with several partners involved in various types of ciliopathies. By integrating proteomic data and functional annotations, we identified a high confidence, ciliary-centrosomal specific LC8 partner list of 57 proteins, 15 of which are central to centriole life-cycle organization. Validated binding motifs and the detailed characterization of the interaction with protein OFD1 emphasized the important role of LC8, which was confirmed by confocal microscopy. OFD1, which is a central player in this system, also stood out as an early and highly conserved LC8 partner. However, additional partners showed a more recent evolutionary origin featuring novel proteins as well as novel motifs. Altogether, our results highlight that LC8 plays a major role in the ciliary-centrosomal system and its interaction network underwent a major expansion. This system driven motif evolution contributed to the increased complexity of the organization and regulation of the ciliary-centrosomal system.

## Introduction

Protein interaction networks are centered around hub proteins that form interactions with a large number of partners. At a molecular level, hub proteins often correspond to specific globular domains that recognize their partners through a common short linear motif (SLiM) [1]. Typically, SLiMs are less than ten residues in length and occur within intrinsically disordered regions. Due to their small size and highly flexible nature, SLiMs generally form transient, context-dependent interactions that play important roles in many regulatory and signaling processes [2]. They can be involved in trafficking and targeting proteins to specific cellular locations, controlling their activities and complex formations and regulating their synthesis and degradation. While in many cases the binding domain presents a highly conserved interface, SLiMs commonly show a high evolutionary plasticity where even a few mutations can lead to the creation or elimination of the interaction site [3]. As a result, SLiMs play essential roles in rewiring the interaction networks depending on tissue specificity and cellular states, and contribute to the evolution of cellular complexity and robustness [4–6]. However, the organizing principles of how changes of SLIM networks are linked to the system-level evolution is largely unexplored.

In the current work, we focus on the SLiM based interaction network of the eukaryotic hub protein LC8, a dimeric protein that has two parallel binding grooves that are recognized by partner proteins [7]. This interface is highly conserved in animal species, remaining practically unchanged from eumetazoans to mammals [7]. In humans, there are two paralogs which share 93% sequence identity. LC8 was initially suggested to function as a cargo adaptor based on its association with the dynein motor complex and its binding to the intermediate chain [7,8]. However, the growing number of partners with a tendency to dimerize lead to the suggestion that LC8 functions as a dimerization hub in a variety of systems [8,9]. LC8 was also shown to drive higher-order oligomerization of already formed dimers, for example in the case of the ciliary protein LCA5 [10]. Until today, more than 100 LC8 binding partners have been experimentally verified [7,11]. These proteins are associated with a diverse set of functions including transport processes, tumor suppression, DNA damage repair, apoptosis, mitosis and regulation, indicating that LC8 has a more general function with a widespread role in the cell [7,12].

Most known partners of LC8 contain a core TQT sequence within the binding region. In order to explore the consensus binding motif of LC8, Rapali et al. used phage display and identified the hyperbinder sequence [12]. LC8 was also subject of a proteomic phage display study using an unbiased peptide library of disordered regions from the human proteome [11]. While both studies identified several new partners, they still provided an incomplete picture of the interactome of LC8. Using an alternative strategy, we have shown that LC8 also participates in the upstream regulation of the Hippo pathway through interacting with AMOT and WWC family members [13]. This approach was based on the known binding partners, and used a powerful bioinformatic pipeline in combination with experimental methods in order to identify and characterize biologically relevant binding motifs. Nevertheless, there were indications of multiple additional partners related to the centrosome. LC8 was shown to be involved in the recruitment of CDK5RAP2 to the centrosome [14]. Anastral spindle-2 (Ana2), a centriole duplication factor specific to Drosophila, was also identified as a binding partner [15]. However, the role of LC8 in the centrosome was not analyzed systematically.

Here, we employed a machine-learning based computational motif prediction pipeline to expand the interaction network of LC8. Similar strategies have been applied to other linear motif binding systems as well [13,16]. We also integrated additional tools into the motif identification pipeline, such as GO enrichment analysis, evolutionary conservation calculation together with the result of mass-spectrometry based interactomics in order to generate a high-confidence binding motif hit list, which was used as a starting point for further in-depth analysis. Our results highlighted the importance of LC8 in the ciliary-centrosome system. We experimentally characterized the interaction of OFD1 (Centriole and centriolar satellite protein OFD1), which, based on its strong evolutionary conservation, could serve as an entry point for LC8 in this system. We also show that this interaction network significantly expanded during evolution, especially in functions related to the life-cycle of centrioles. Our study showcases a striking example of the co-evolution of the SLiM-based interaction network with cellular processes.

## Results

### New instances of known LC8 binding motifs identified based on machine learning

In order to identify new LC8 binding instances in the human proteome, we trained a machine learning method to capture the specific sequence features that are shared among linear motif sites that bind to LC8. As a starting point, we collected a set of 91 experimentally verified human LC8 binding motifs (Supplementary Materials). Given the limited number of examples, we chose to implement a random forest classifier, as tree-based methods were shown to outperform more complex neural networks on small datasets [17]. For the classifier, we gathered various features which can be associated with the experimentally verified LC8 binding motifs. The common sequence motif was described using an eight amino acid length position specific scoring matrix (PSSM). As expected, the PSSM clearly describes the highly conserved TQT central tripeptide, assigning the highest scores for these amino acids (fig. S1). Additional biochemical and protein structure based features were also incorporated into the feature set. Disordered region prediction was calculated by applying the biophysics based IUPred algorithm [18]. Structure based disorder information was integrated by calculating pLDDT values and relative solvent accessibility (RSA) from the pre-built AlphaFold2 models [19,20]. Disordered binding site prediction was added to the features by using the ANCHOR2 tool [21]. Protein domain information gathered from Pfam was also used as a feature by identifying positions with overlap of Pfam domain annotations [22]. Secondary structure prediction was not used in the feature set as it has been shown in our previous work that this criterion did not have a strong discriminatory power for the LC8 system [13]. In total, we trained our classifier on 6 features by using the experimentally verified human instances as a positive set and 10000 random human sequences with the same length as a negative set.

Stratified five-fold cross-validation produced an average accuracy score of 0.9904 with a balanced accuracy of 0.842. The average true positive rate was 0.69, which resulted in the correct categorization of 63 instances from the 91 experimentally validated human binding motifs by summing over all five possible test sets. The false positive rate was very low, averaging 0.007. The cross-validation procedure yielded an average area under the Receiver Operating Characteristic (ROC) curve of 0.967 with standard deviation of 0.0185 (Fig. 1A) (Supplementary Materials). Relative importance scores for each input feature of the trained model were calculated based on the reduction of the entropy criterion used to select the split points. As expected, the strongest criterion was the PSSM feature with a score of 0.772. The second criterion was the pLDDT with a score of 0.114. The third, fourth and fifth criterion were RSA, IUPRED and ANCHOR with less than 0.1 scores (0.06, 0.035 and 0.019, respectively). The Pfam feature contributed minimally to the classifier (0.001) (Fig. 1B).

**Fig. 1.**
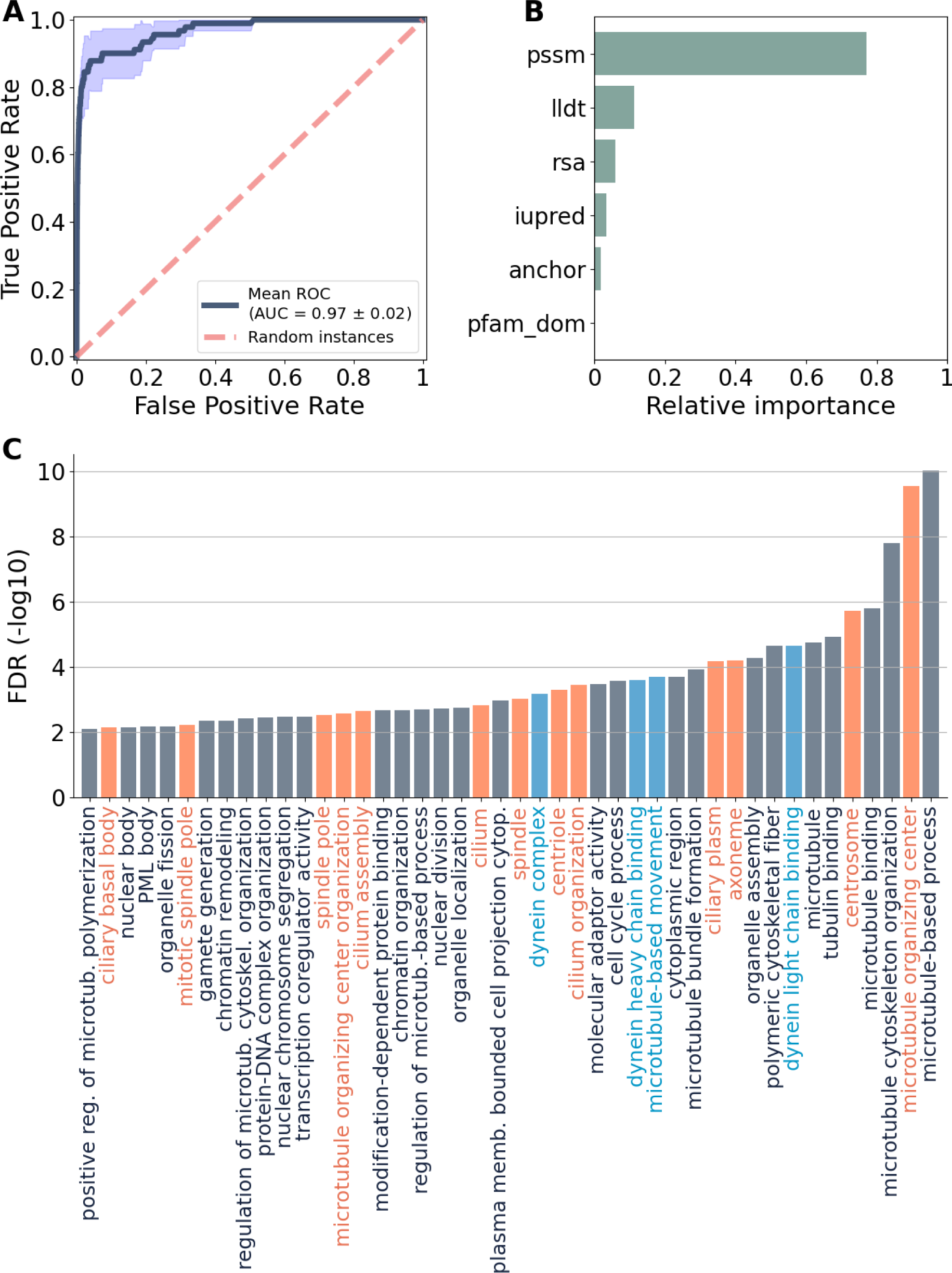
Cross validation and relative feature importance of the random forest model, and significantly enriched GO terms associated with LC8 partners. **(A)** Stratified five-fold cross-validation ROC curve. **(B)** The calculated relative feature importance scores of the trained classifier. **(C)** False Discovery Rate (FDR) of the significantly enriched, LC8 partner associated terms (all three GO biological aspects in one, for details see supplementary-materials). Ciliary-centrosome system and the dynein complex associated terms are indicated with orange and blue, respectively.

Using the trained classifier, we scanned the human proteome for novel binding motifs. For this, we considered every 8 amino acid long peptides of the human proteome after filtering out secreted proteins and extracellular/transmembrane protein regions. The confidence score for all instances was calculated based on the evaluation of the random forest classifier. Our model identified 896 novel instances as putative binding motifs in 768 proteins. The number of predicted novel instances was narrowed down by filtering out motifs with confidence scores below that of the lowest (0.598) scoring known binding motif (_529_KQAYVQCN from DYNC2I1) (Supplementary Materials), which resulted in 725 hits of 628 proteins. Based on this filtered motif hit list together with the 91 instances of the 67 known partners we highlight our LC8 binding motif list composed of 816 instances of 674 proteins. Among the 674 partners, interaction data with LC8 collected from the IntAct and Biogrid protein-protein interaction databases was available for 139 proteins of the 674 LC8 partners, in total (for 92 predicted and 47 known partners) [23,24].

### LC8 partners are enriched in the ciliary and centrosomal systems

Next, using the 674 LC8 partners, we carried out a functional enrichment analysis based on Gene Ontology (GO) to determine biological functions over-represented among LC8 binding proteins. The GO term over-representation test against the human proteome was performed on the GOC website using the enrichment analysis tool from the PANTHER Classification System [25–27]. A total of 44 significantly enriched terms were obtained, 7, 20 and 17 Molecular Function, Biological Process and Cellular Component GO aspects, respectively (Supplementary Materials). As expected, the dynein motor complex associated terms (Fig. 1C) were significantly enriched with 11 LC8 partners, in total. Eight of these proteins are the well-known dynein chains, including four cytoplasmic dynein intermediate chains (DYNC1I1/2, DYNC2I1/2) responsible for transport activity, and four dynein axonemal intermediate (DNAI1/2/3/4) associated with motile cilia movement. Out of the 44 significantly enriched terms, a total of 13 terms (30%), such as centrosome, ciliary basal body and cilium, were tightly associated with the ciliary-centrosome system (Fig. 1C). There were a total of 115 LC8 partners, including well-studied proteins, such as CDK5RAP2, LCA5, CEP152 and CENPJ, that were associated with these 13 ciliary-centrosome system related GO terms. The vast majority of the 115 LC8 partners, 87 proteins in total, were linked to the two major cellular component terms of the ciliary-centrosome system, the Cilium and Centrosome (Supplementary Materials).

Using the enrichment tool g:Profiler [28], we also noticed that the 674 LC8 partner proteins showed a significant enrichment (negative logarithm of the P-value is 5.8) with various types of ciliopathies. Ciliopathies are rare inherited developmental disorders that arise from defects of primary or motile cilia structure or function [29]. The heterogeneous phenotypic spectrum of ciliopathies is connected to mutations of approximately 200 genes associated with motile and non-motile cilia [30]. Altogether, we identified 26 potential LC8 binding partners connected to ciliopathy syndromes (Table 1.), based on the data collected by g:Profiler analysis (23 genes) and additional literature survey (3 genes). The unexpectedly high rate (more than 10% of all ciliopathy genes are putative LC8 partners) of ciliopathy genes among potential LC8 partners highlighted the central role of LC8 partners in cilia formation and function. However, according to the ClinVar database [31], pathologic mutations did not directly target the LC8 binding motifs. Most of the mutations were truncation mutations which are likely to lead to the loss of the entire protein. Although we did not find any direct evidence connecting LC8 binding motifs to the development of ciliopathies, results of the enrichment analyses highlighted a new biological aspect of LC8 partners associated with the ciliary-centrosome system.

**Table 1.**
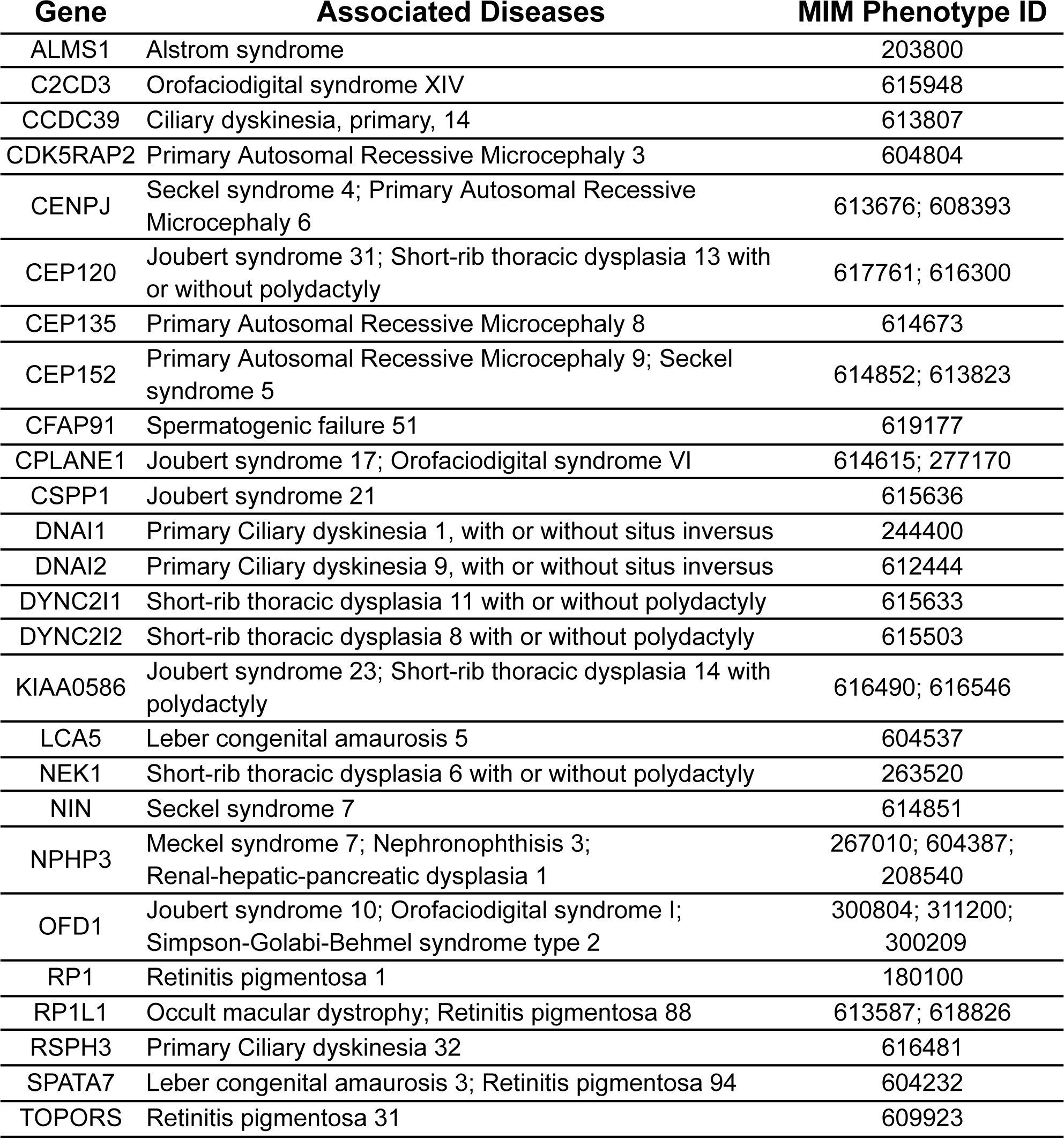
Ciliopathy syndromes associated with predicted LC8 binding partners.

### Integrating high throughput localization and GO annotations show a clear overlap of the ciliary and centrosomal systems

In recent years, a more detailed map of the organization of ciliary and centrosomal systems have started to emerge thanks to various high-throughput proteomic studies, such as mass spectrometry combined pull-down and proximity-dependent biotinylation (BioID) assays [32–40]. Here, we first collected proteins associated with Cilium and/or Centrosome terms using these High-Throughput (HT) studies. This step resulted in 2213 ciliary and 958 centrosomal proteins, in total. Next, we retrieved all proteins from the PANTHER Classification database that had Cilium and/or Centrosome cellular component GO term annotations. We created a ciliary-centrosome protein list based on the intersection of the two sources for both the centrosomal and ciliary protein sets. This yielded 246 centrosomal and 583 ciliary proteins, in total (Fig. 2, A and B). We also collected proteins associated with the dynein machinery by using the GO term Dynein complex, which resulted in 55 proteins, in total (Supplementary Materials). Based on the 884 proteins of the three systems, we established a ciliary-centrosome-dynein protein list (Fig. 2C).

**Fig. 2.**
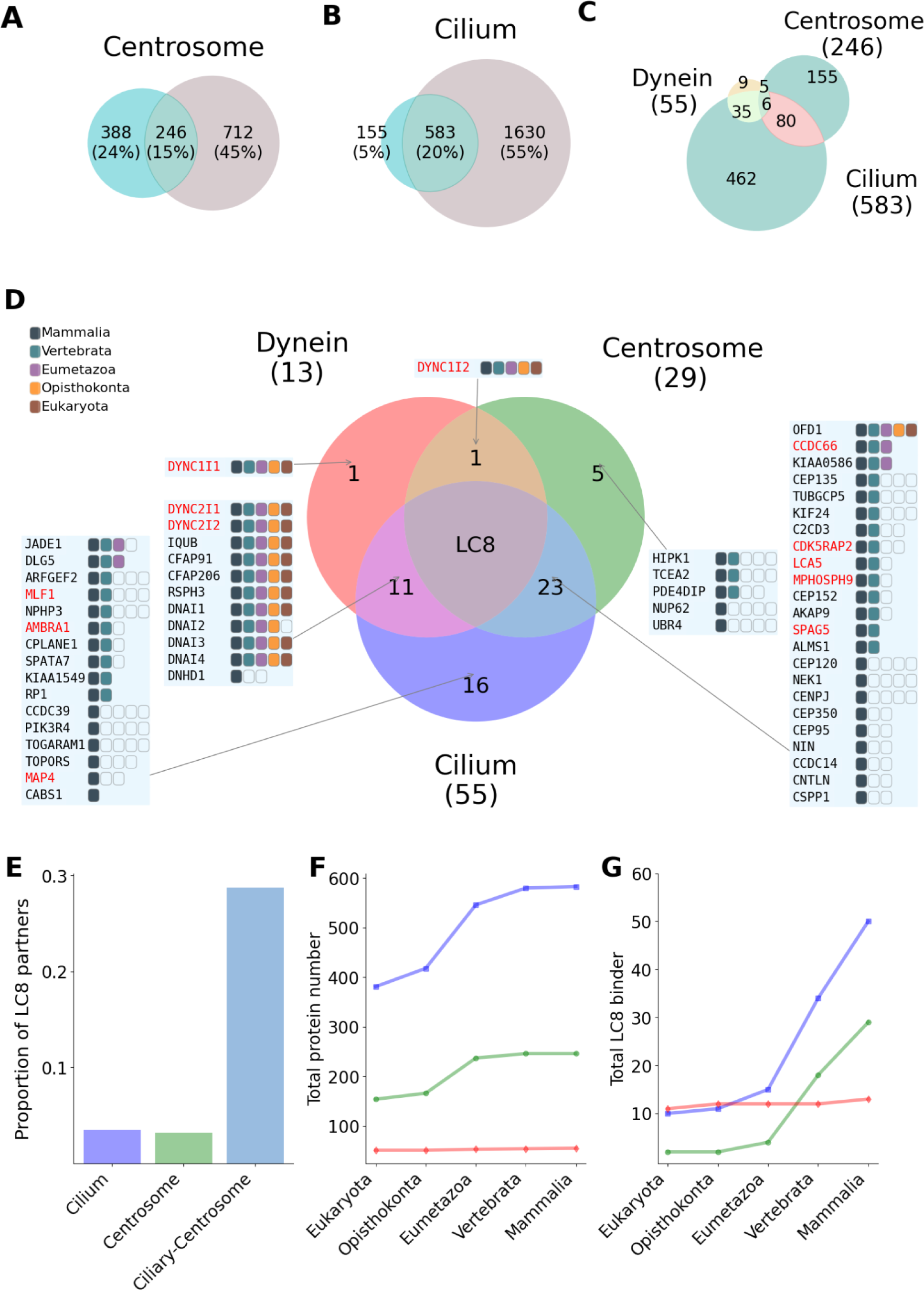
LC8 network in the ciliary-centrosomal system and its evolutionary trace back. **(A-B)** The result of the GO annotation and high throughput (HT) based protein collection of the ciliary and centrosomal systems depicted by Venn diagram. GO and HT sources are indicated by coral blue and light brown, respectively. The number of collected proteins are shown in the circles. Intersections of the two sources are indicated by sea green (ciliary-centrosome protein list). (**C**) The Venn diagram of the protein numbers of the ciliary, centrosomal and dynein systems (ciliary-centrosome-dynein protein list). The exact protein numbers of the three systems are presented in brackets. For the ciliary and centrosomal systems the ciliary-centrosome protein list was used, while for the dynein system the collection of the 55 proteins was used. (**D**) Venn diagram of LC8 partners in the three systems and their evolutionary conservation at both binding motif and protein levels. The conservation of motifs is indicated by colored squares at different evolutionary levels, while the conservation when only the protein is conserved is depicted by empty rectangles. Partners with experimental evidence at the motif level are shown in red color. (**E**) The proportion of LC8 partners among all ciliary and centrosomal proteins. The colors of the three systems correspond to the colors of the Venn diagram of LC8 partners. (**F-G**) Evolutionary expansion of the centrosome, cilium and dynein systems in terms of all proteins (F) and LC8 partners (G) involved in the systems. The evolutionary graphs of the three systems are depicted with green, blue and red, respectively (colors correspond to the colors of the Venn diagram of LC8 partners). Evolutionary expansion of the LC8 network was calculated based on motif conservation.

We found a clear overlap between the ciliary and the centrosomal systems, altogether 80 proteins were associated with both systems (Fig. 2C). They also overlapped with the dynein motor complex, especially with ciliary proteins. In total, there were six proteins involved in all three systems: the two LC8 paralogs (DYNLL1, DYNLL2), DYNLRB1, DYNLRB2, DYNC1H1 and the DCTN1 (Fig. 2C). This further highlights the central role of LC8 in the intersection of the dynein and ciliary-centrosomal systems.

Within the ciliary-centrosome-dynein protein list, 57 proteins contained LC8 binding motifs, from which 30, 54 and 13 proteins were annotated for the centrosome, cilium and dynein systems, respectively (Fig. 2D). These three sets also showed a significant overlap (Fig. 2D). The majority of proteins with LC8 motif within the dynein motor complex (11 out of 13) were also associated with the cilium. The intersection of the dynein motor complex and the centrosome contained a single protein (DYNC1I2), while there was only one member of the dynein complex that was not associated with neither the cilia nor the centrosome (DYNC1I1). 16 proteins were associated only with the cilium and 5 had only centrosomal annotations. The largest group in this network was formed by the intersection of cilium and centrosome, involving 23 proteins (Fig. 2D).

Comparing the number of LC8 partners in the ciliary-centrosomal (CC) system with the total number of proteins in the system, we found that 28.8% (23:80) of the CC system proteins were LC8 binders. This is a higher ratio compared to if the ratios for cilium and centrosome are calculated separately. In these cases, the proportions of LC8 partners are only 3.5% (16:462) and 3.2% (5:155) for the cilium and centrosome systems, respectively (Fig. 2E). These results clearly demonstrate the central role of LC8 in the ciliary-centrosomal system.

### Emergence of LC8 binding partners

We also analyzed the evolutionary emergence of proteins and their binding motifs of dynein, cilia and centrosomal systems. We traced back the conservation of these proteins through orthologous sequences grouped at five evolutionary levels which corresponded to the major taxa of Mammalia, Vertebrata, Eumetazoa, Opisthokonta, Eukaryota.

At the protein level, nearly all components of the dynein complex showed an ancient evolutionary origin. The ciliary and the centrosomal systems also contained several highly conserved proteins which could be traced back to the common ancestors of eukaryotes. However, these two systems accrued many more new members during evolution. The major expansion happened before the emergence of eumetazoans. The rates of expansion between eukaryotic and mammalian levels were 1.5-, 1.6- and 1.1-fold for the ciliary, centrosomal and dynein systems, respectively (Fig. 2F). These results are consistent with previous knowledge that the dynein complex is a more conserved functional unit, while the ciliary-centrosomal system is a dynamically changing system [41].

We also evaluated the conservation of the binding motifs. For this, we checked if presence of a binding motif could be detected in the majority of orthologs at a given taxa level and based on the most distantly related level determined the evolutionary emergence of the motif (Fig. 2, D and G). In the dynein complex, the majority of motifs showed an ancient origin with Eukaryota level conservation, with the only exception of the DNHD1 motif which appeared at the mammalian level. In contrast, the LC8 interaction network underwent a dramatic expansion within the centrosome and cilia systems, which started at the level of Eumetazoa, but continued to extend even at the levels of Vertebrate and Mammalia. In the ciliary and centrosomal systems 10 and 2 ancient, eukaryotic motifs were identified, while at the mammalian level the numbers of conserved motifs were 50 and 29, respectively, which represent 5 and 14.5-fold increases, respectively. In the dynein system the expansion of LC8 partners was only 1.2-fold (11 to 13) (Fig. 2G). Altogether, our results demonstrate that LC8 interacts with ancient members of the ciliary-centrosome system but it gained multiple new partners at multiple steps during evolution to which both the emergence of novel proteins and the emergence of novel motifs in already existing proteins contributed.

### OFD1 interacts with LC8 on the protein level through a short linear motif

Our results highlighted OFD1 as the most conserved non-motor function related LC8 partner in the human proteome. OFD1 plays a critical role in the centriole organization and is tightly connected to ciliogenesis [42,43]. OFD1 was previously shown to interact with LC8 using tandem affinity purification coupled mass spectrometry [44]. Binding peptide of OFD1 was also identified using phage display, but the thermodinamic properties of the binding and the dissociation constant have not been published [11]. Here, we further characterize the interaction between OFD1 and LC8.

The LC8 binding motif of OFD1 is located at the N-terminal (_157_CNMETQTS_164_), between a dimerization LisH domain and an extensive coiled coil region (Fig. 3A). The motif is highly conserved across eukaryotic lineages and could be traced back to unicellular algae *C. reinhardtii*, which suggests that an ancient binding motif could be present in the common ancestral of humans and algae (Fig. 3B). Among unicellular organisms, orthologous binding motifs in yeast such as *S. cerevisiae* were not detected, in agreement with previous results that yeasts have a centriole-less centrosome [45]. Using a short synthetic peptide of the binding motif (KESCNMETQTS), the dissociation constant between the OFD1 peptide and LC8 was measured by competitive fluorescence polarization assay against the FITC-labeled known LC8 binding motif of BMF as 12.8 ± 1.7 μM (Fig. 3C). We also validated our result with SPR method which resulted in a similar K_d_ (12.5 ± 1.3 μM) (fig. S2). To complement motif level analysis, we characterized the interaction OFD1 with LC8 at the protein level. Similarly to many large coiled coil containing centrosomal proteins, OFD1 is poorly soluble and tends to aggregate in solution. Therefore, we used a truncated form of OFD1 containing a 270 residue long N-terminal construct (sOFD1-WT) and fused an MBP tag to further increase its solubility (fig. S3). We also designed a motif mutant form (sOFD1-ΔTQT) in which the central TQT core of the motif was substituted with alanines (fig. S3). For the protein level validation, we performed a GST pull-down experiment between GST-LC8 and the sOFD1 constructs. The sOFD1-WT showed interaction with GST-LC8, but the introduced Ala mutations in the sOFD1-ΔTQT abolished the binding between the two proteins (Fig. 3D). These results indicated that the LC8 could bind to OFD1 at the protein level, and the interaction was mediated by the previously identified and validated linear motif.

**Fig. 3.**
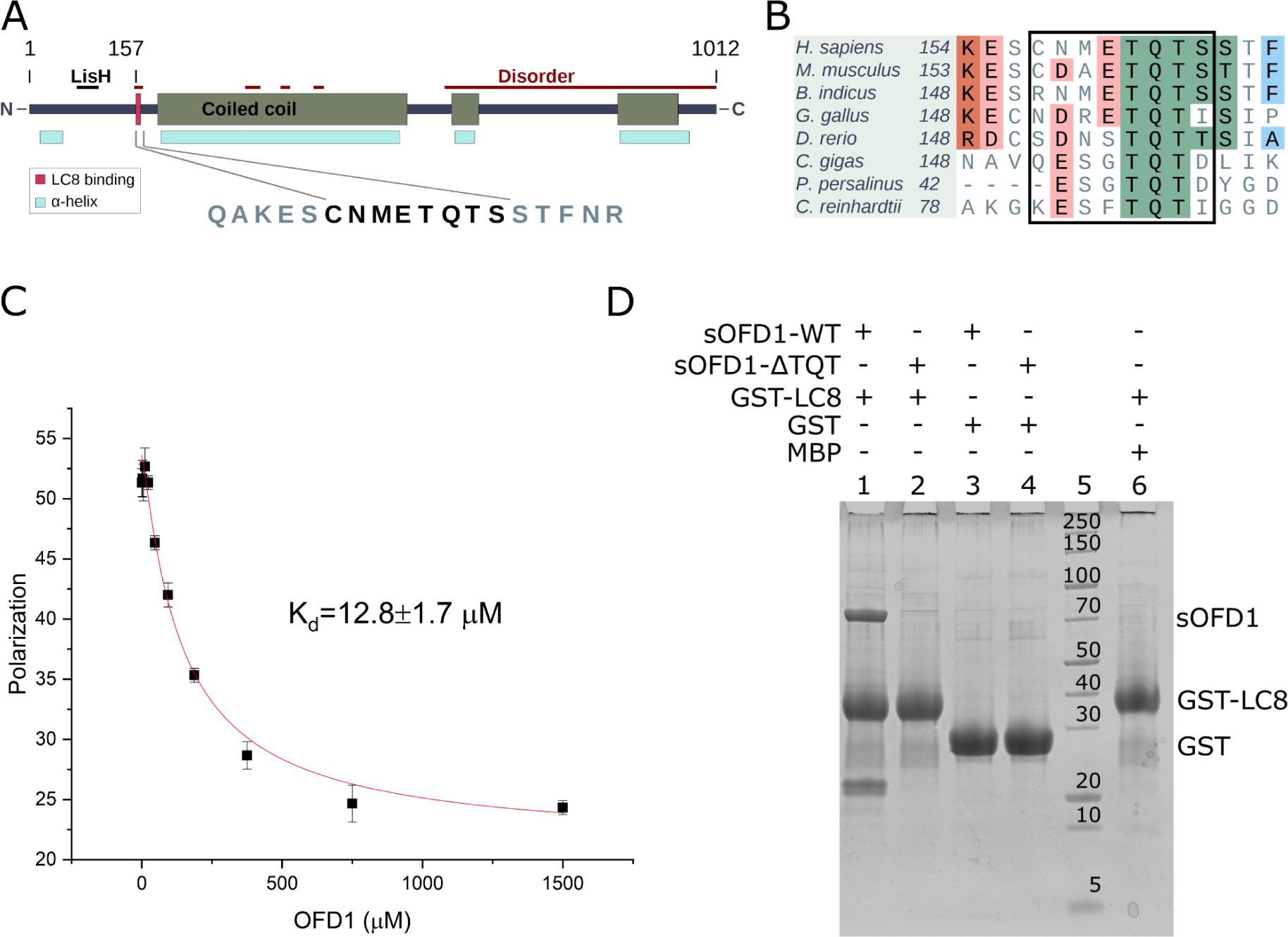
Sequence localization and conservation of the LC8 binding motif of OFD1, and the peptide and protein level validation of the OFD1-LC8 interaction. (**A**) The domain structure of the full-length OFD1 protein. (**B**) The multiple sequence alignment of the sequence region of LC8 binding motif generated from the corresponding OFD1 orthologs of model organisms. The 8 residue long motif is indicated by a rectangle. (**C**) The dissociation constant (including SE) of the LC8-OFD1 interaction was measured in triplicate (n = 3) by a competitive fluorescence polarization assay. (**D**) The GST-pull down assay between GST-LC8, sOFD1-WT, and sOFD1-ΔTQT and the control experiments using sOFD1 constructs with GST and GST-LC8 with MBP.

### LC8 colocalizes with OFD1 in the centrosome and the basal body

According to literature data, OFD1 is a characteristic centriolar satellite protein. In interphase cells, OFD1 is mostly localized in the pericentriolar satellites [43] but is also present at the centrosome [42]. During cilia formation and in the presence of mature primary cilia OFD1 is localized in the basal body [43,46]. Our previous results showed that LC8 is localized mainly in the ciliary basal body and in the centrosome [10]. To evaluate the possible localization of the LC8-OFD1 interaction in ciliated and non-ciliated hTERT-RP1 cells we performed a confocal microscopy study.

In ciliated cells OFD1 was localized in the basal body and indicated a minor entry into the ciliary axoneme. The colocalization with LC8 was observed in the ciliary basal body (Fig. 4A). The mean pixel intensity based correlation between the LC8 and OFD1 was 0.607 ± 0.028 (Pearson’s coefficient) and the overlap characterized by the thresholded Manders coefficient (tM) related to the OFD1 pixels was tM(OFD1) = 0.481 ± 0.024 and tM(LC8) = 0.527 ± 0.023 for LC8.

**Fig. 4.**
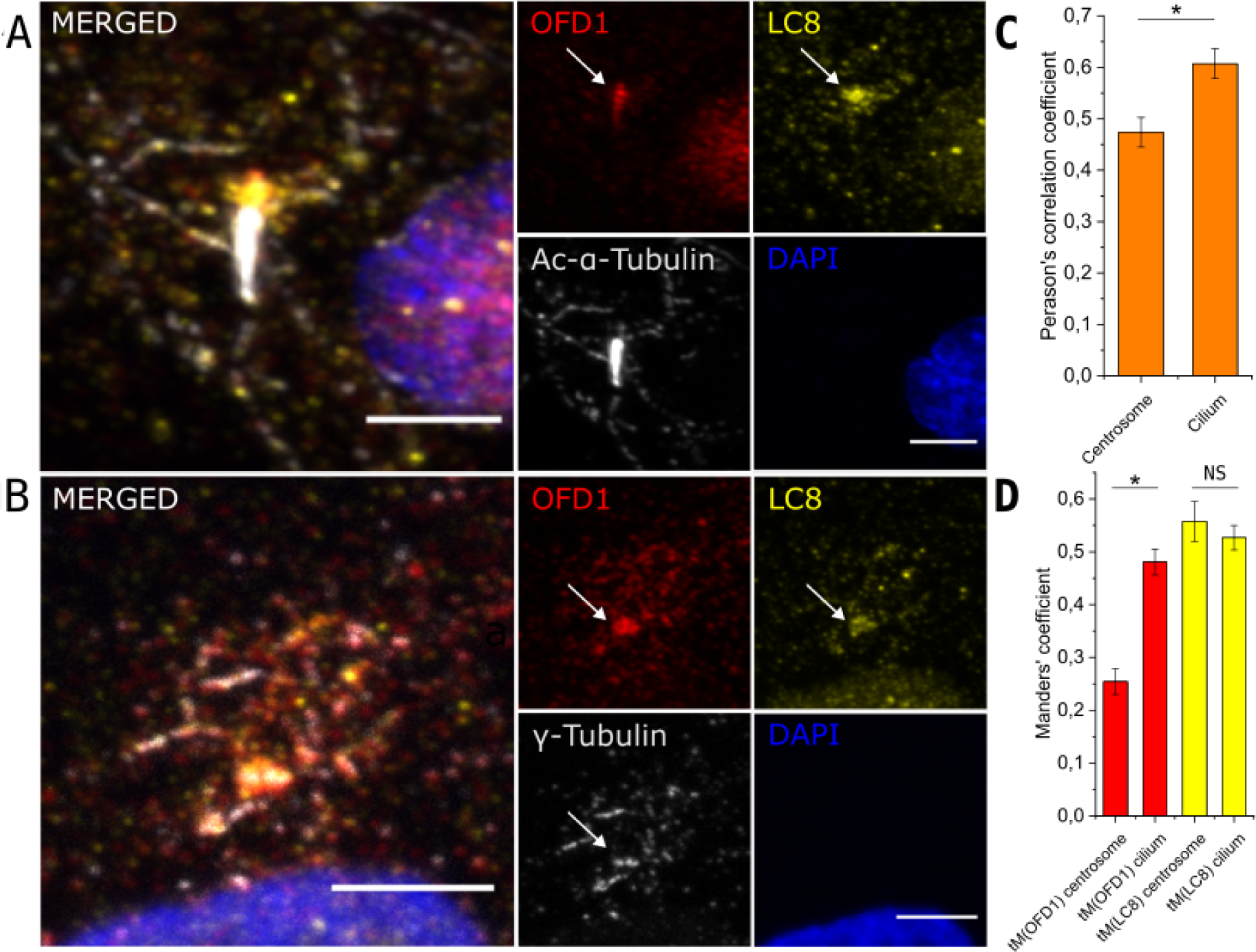
The localization of LC8 and OFD1 in ciliated and non-ciliated cells. (**A**) LC8 (yellow) and OFD1 (red) localized in the ciliary basal body (indicated by white arrows) of ciliated hTERT-RPE1 cells. OFD1 also entered the ciliary axoneme stained by acetylated α-tubulin (white). The merged image shows the colocalization of the two proteins in the ciliary basal body. Scale bar, 4 μm. (**B**) LC8 (yellow) and OFD1 (red) localized in the centrosome of non-ciliated hTERT-RPE1 cells overlapped with γ-tubulin (white) indicated by white arrows. The colocalization of LC8 and OFD1 is shown in the merged image. Scale bar, 4 μm (**C**) The pixel intensity based on Pearson’s correlation coefficient comparison of the LC8 and OFD1 in the centrosome and the ciliary basal body. (**D**) The overlap of OFD1 and LC8 pixels and LC8 and OFD1 pixels, respectively in the centrosome and the ciliary basal body. Error bars in (C) and (D) represent SEM; n = 7. NS, not significant (P ≥ 0.05); *P < 0.05, Student’s unpaired two-sample t-test.

In non-ciliated cells, OFD1 was detected in the centrosome and around the centrosome in the pericentriolar satellites. LC8 was also present in the centrosome and overlapped with γ-tubulin. However, it was barely detectable in the pericentriolar satellites (Fig. 4B). The mean of the calculated Pearson’s correlation coefficients of LC8 and OFD1 in non-ciliated cells was 0.474 ± 0.028 which was significantly lower compared to the measured value of the ciliated cells (0.607±0.028) (Fig. 4C). The mean of the Manders coefficients of the OFD1 (tM(OFD1) = 0.254 ± 0.024) was also significantly lower in non ciliated cells, but no significant difference was detected in case of the LC8 (tM(LC8) = 0.557 ± 0.037) (Fig. 4D). This could be explained based on the observation that OFD1 is also localized in the pericentriolar satellites in non ciliated cells, but it is removed by autophagy from the satellites during cilia formation [43] whereas LC8 is restricted to the centrosome and the basal body. Altogether, these results showed that the interaction between LC8 and OFD1 is mostly confined to the centrosome and to the basal body, despite the strong presence of the OFD1 in the pericentriolar satellites and in appearance ciliary axoneme.

### LC8 plays central role in the centriole lifecycle organization

To better understand the role of LC8, we drew a more detailed landscape of the ciliary-centrosomal interactome based on annotations and literature data (Supplementary Materials). We divided the 57 partners into 14 subgroups (Fig. 5) highlighting the detailed organization of this system. The central role of LC8 partners in the ciliary-centrosomal system is also emphasized by their overrepresentation (25 out of 57) in ciliopathy related genes (Table 1). We further validated seven new LC8 binding partners, which represent some of the notable features, such as disease association or evolutionary conservation (Table 2). LC8 partners already known or validated in this study are highlighted on Fig 5.

**Fig. 5.**
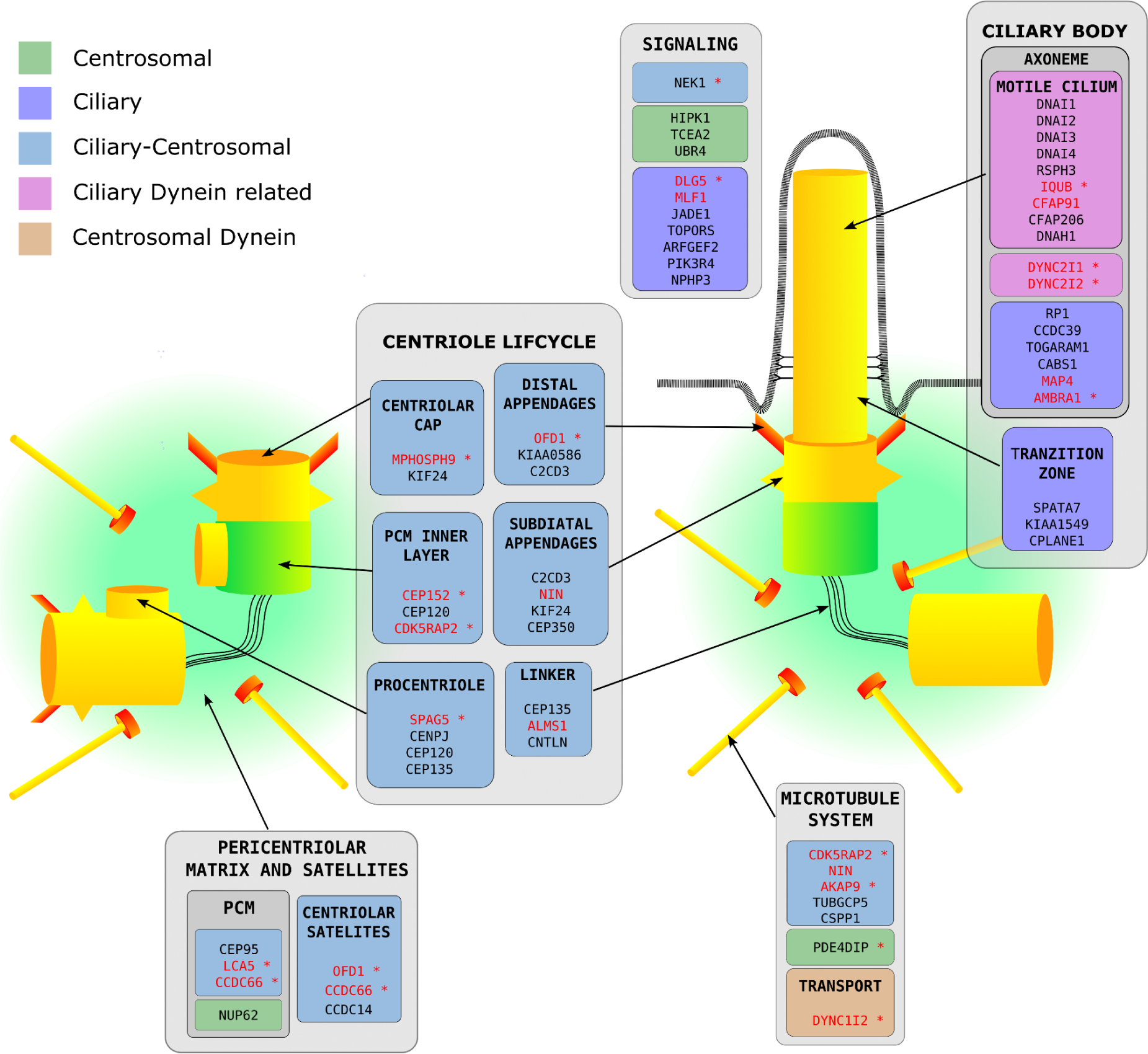
The localization and functional associations of the predicted LC8 binding partners. On the schematic representation of the centrosome and the primary cilia, the main anatomical features and components of the system were represented, and associated with colored text boxes. The colored text boxes grouped together in light gray boxes represent the major aspects that are associated with LC8 functions. The text boxes contain the name of the corresponding component and the associated LC8 binding partners. The text boxes were colored according to the association of the LC8 binding partners to the centrosome (green), cilia group (dark blue), both cilia and centrosome (light blue), ciliary dynein related partners (purple) and the centrosomal dyneins (brown). The proteins with known LC8 binding motifs (including the ones validated in this study are highlighted in red, the additional red asterisk after the protein names means HT data indicates the interaction with LC8 according to the BioGRID and IntAct databases [23,54].

**Table 2.** LC8 binding motifs validated in partners associated with ciliary centrosomal functions. The dissociation constants (K_d_) are measured by SPR technique (IQUB, MAP4, CFAP91, CEP152) and fluorescence polarization (CDK5RAP2, AKAP9, ALMS1) in triplicates (n = 3)

| Gene Name | UniProt ID | START | Motif Sequence | K<sub>d</sub>(μM)±SE(μM) |
| --- | --- | --- | --- | --- |
| IQUB | Q8NA54 | 266 | HNAGTQTV | 3.57±0.13 |
| MAP4 | P27816 | 800 | GSKSTQTV | 4.42±0.68 |
| CFAP91 | Q7Z4T9 | 196 | RDADVQTD | 0.55±0.03 |
| CEP152 | 094986 | 789 | LDCGSQTD | 2.37±0.29 |
| CDK5RAP2 | Q96SN8 | 869 | KDASVQTV | 12.46±3.88 |
| AKAP9 | Q99996 | 1679 | RNSSTQTQ | 12.40±3.08 |
| ALMS1 | Q8TCU4 | 2334 | KDIGTQTN | 19.20±5.50 |

One of the major groups of LC8 partners is associated with the ciliary body, harboring axonemal and transition zone related proteins. (Fig 5). These proteins are ancient dynein related partners, associated with the intraflagellar transport (DYNC2I1, DYNC2I2) and the force-generating machinery of the motile cilia, including the inner and outer dynein arms (DNAI1,2,3,4). Several partners are associated with the radial spoke, including IQUB and CFAP91, validated at the motif level here (Table 2; fig. S2), as well as CFAP206, RSPH3. Additional cilia specific partners, such as MAP4 which was also validated here (Table 2), are connected to the microtubule system of the axoneme, the transition zone, or are involved in photoreceptor functions (Supplementary Materials).

Another major group represents the intersection of ciliary and centrosomal proteins with 15 partners (Supplementary Materials). These proteins are tightly connected to the lifecycle of the centriole, further emphasizing the central role of LC8 in this system. The putative partners connect LC8 to the procentriole formation, to the ciliary cap, which is one of the main suppressor of ciliogenesis, and to the core components of the inner layer of the pericentriolar matrix (PCM). They are also associated with the distal and subdistal appendages involved in centriole maturation and ciliogenesis, and the linker complex responsible for centriole cohesion (Fig. 5). These putative LC8 binding partners also interact with each other and form subnetworks, or recruit each other to specific cellular features [47–50]. We validated three additional LC8 binders in this system, including the PCM inner layer related CEP152 [51], the already identified but kinetically not characterized CDK5RAP2 [14] and the linker-associated ALMS1 [52].

Beyond the centriole organization and the ciliary functions, LC8 also participates in the organization, and the nucleation of the microtubule system. (Fig. 5) In this system, we identified and validated the AKAP9 protein (Table 2), which is an anchoring component of the γ-tubulin ring complex [53].

LC8 is also associated with PCM related proteins and partners associated with the pericentriolar satellites. The satellite associated LC8 partners are also presented in the PCM (CCDC66) and the distal appendages (OFD1) (Fig. 5, Supplementary Materials). According to our confocal microscopy results, the interaction between satellite related proteins and LC8 might take place in the PCM and in the centrioles instead of in the satellites.

The ciliary centrosomal system is also a signaling hub. In agreement, LC8 partners also contain elements of the WNT and Hedgehog signaling pathways and signaling routes associated with ciliogenesis and cellular stress response (Fig. 5, Supplementary Materials).

Altogether, our results demonstrate that LC8 partners are connected to every aspect of the ciliary-centrosomal network, suggesting that LC8 plays a critical role in the detailed organization of the ciliary-centrosomal network. While its connection to dynein related functions is highly conserved across a wide range of eukaryotic species, however, its participation in the centriole lifecycle organization with the exception of OFD1 is a more recent evolutionary origin.

## Discussions

LC8 is a eukaryotic hub protein that recognizes its partners through a short linear motifs binding to a highly conserved interface. In the last few decades, over a hundred LC8 binding motifs were identified and validated in a functionally diverse set of binding partners [7,11,13]. However, the comprehensive functional analysis of the interaction network of LC8 was still incomplete. In this work we applied a machine learning approach to generate an upgraded high-confidence partner list and identified 628 putative LC8 binding partners in the human proteome. We carried out an in-depth analysis of functional enrichment combined with evolutionary properties in order to gain a better understanding of the LC8 interaction network at the system level. In agreement with previous results, we found that LC8 was strongly associated with the dynein-related motor complexes of the intraflagellar transport and the force-generating machinery of the motile cilia [55–58]. However, the novel partner pool, which combined the putative binding motifs with the known instances in the human proteome, also highlighted that LC8 plays a significant role in the ciliary-centrosomal system. This was further emphasized by the association of multiple partners with various types of ciliopathies.

Previous proteomic studies have shown that the cilia and the centrosome form overlapping interaction networks centered around the centrioles [36,59]. In dividing cells centrioles maintain the complex network of the pericentriolar matrix proteins to orchestrate the functions of the centrosome. In non-dividing cells they form the ciliary basal body to organize the cilia formation and protein trafficking of the ciliary body. During the cell cycle, the centrosomes go through a precisely controlled lifecycle organized by the associated proteins of the centrosome and the basal body [35,60,61]. By integrating high throughput data we identified putative LC8 binding motifs in 57 proteins associated with discrete ciliary and centrosomal functions. The most pronounced enrichment of LC8 binding partners was found in the intersection of the centrosome and cilia. At a closer look, we found that LC8 partners are major players in the centriole lifecycle organization and are tightly connected to centriole integrity, duplication, and maturation related processes. The identified partners are also involved in the nucleation of the cellular microtubule system, the organization of the centrosomal and ciliary structure, orchestrating the ciliogenesis and they participate in the centrosome and primary cilia related signal transduction. LC8 partners are also found in the pericentriolar matrix of the basal body and the centrosome, in the centriolar satellites, and in the ciliary transition zone. Several of the predicted partners are verified at the motif level and in the case of OFD1, which is a central player in this system, also at the protein level. The localization pattern of LC8 obtained by confocal microscopy also confirmed the involvement of LC8 in this system.

It was suggested that the centrosome evolved from the basal body/axoneme of the unicellular ancestor of eukaryote, preserving the association of three basic cellular functions of sensation, motion and division [41]. While the dynein motor complex has shown limited diversification across major evolutionary transitions, the centrosome had a highly complex evolutionary history with major components being lost and reinvented at multiple times [45]. In agreement with these different scenarios, LC8 partners associated directly with the dynein motor complex generally showed ancient evolutionary origin which could be traced back to early unicellular eukaryotic lineages. In contrast, among the non-dynein related LC8 partners, only OFD1 showed similar evolutionary conservation. This further emphasized the central role of OFD1 in the centrosomal interaction network. However, the other LC8 binding motifs associated with the ciliary-centrosomal system were more recent evolutionary inventions. The LC8 network underwent a major expansion at multiple stages of evolution involving both the appearance of novel proteins and novel motifs within already existing proteins. Instead of the random occurrence and disappearance of short linear motifs due to a few mutations [3], we observed here a system level expansion of the motif network co-evolving with the highly complex centrosome-ciliary system.

An interesting question remains, what is the precise role of LC8 in the centrosome. The centrosome was suggested to form a biomolecular condensate probably with liquid-like properties [62,63]. The formation of such membraneless organelles is driven by multivalent weak interactions. In the case of the centrosome, coiled coil like helical segments are likely to play such a critical role. Many of the putative partner proteins of LC8 contain coiled coil segments, especially in the set associated with the centriole. While LC8 was originally suggested to drive the dimerization of its binding partners, LC8 was reported to promote the higher-order oligomerization of proteins, for example in the case of ANA2, NEK9, and LCA5 [10,15,64]. This can lead to an interesting hypothesis that LC8 plays an important role in the structural organization of membraneless organelle of the centrosome. However, the elucidation of the precise role of LC8 within this system requires further studies.

## Materials and Methods

### Random Forest based computational pipeline

#### Collecting experimentally validated LC8 binding motifs

We created a manually curated human protein list of LC8 binding partners based on literature search, in which the interaction with LC8 was verified at the binding motif level. The length of the core binding motif was taken as 8 amino acids based on the structure of LC8 and DYNC1I2 (PDB:2PG1), in which the 8 amino acids of the binding region form contacts with LC8. The final database contained 91 motif instances of 68 partners. From the 91 known binding motifs, 49 contained the canonical TQT motif core. The verified motifs collected from the literature and their sources are summarized in Supplementary Materials.

#### Sequence features

For the random forest based model we considered six features for the training model, which were the known instances based Position Specific Scoring Matrix (PSSM) and five further biochemical and structural features, disorder prediction results of IUPred, ANCHOR and AlphaFold2, and motif overlap with Pfam domain predictions.

To generate an LC8 binding motif specific PSSM, we collected a total of 91 human binding motifs from the literature for which there was experimental evidence (Supplementary Materials). The motifs were applied for PSSM constructing as 8 long regions in which positions 5, 6 and 7 were the consensus TQT tripeptides. The elements of the PSSM (P_i, j_) were expressed as the log-odds score of amino acid frequency in each position in the known motifs divided by the background frequency. As not every amino acid was present in each position in the known set, a pseudo-count correction was introduced.

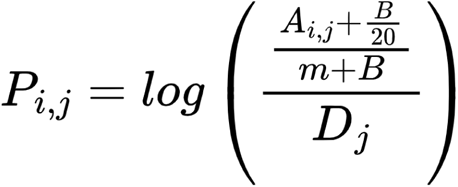

Where A_i, j_ is the frequency of amino acid i at position j in the known motif set, and D_i_ is the background frequency of amino acid i. The background frequency was calculated using the eukaryotic proteome from UniProt. B is the pseudocount with a value of 5 [65], m is the number of sequences, and 20 is the number of amino acids.

The pre-calculated AF2 models for human proteins were collected from the FTP site (https://ftp.ebi.ac.uk/pub/databases/alphafold) (version 4). This dataset contained compressed PDB files for 20504 human proteins, including proteins longer than 2700 amino acids that were provided as 1400 residue long, overlapping fragments. Residue specific pLDDT and RSA (relative solvent accessibility) scores of proteins were calculated from the downloaded PDB files using the AlphaFold-disorder python tool, which parses and processes AF2 based PDB files and returns pLDDT scores based on the B-factor of structure and calculates RSA as provided by DSSP and BioPython [66,67]. pLDDT and RSA for each instance were calculated as the mean value of the instance match positions.

For disorder and disorder binding prediction, the most recent version of IUPred and ANCHOR algorithms were employed with default parameters [18]. Information of protein domains was generated by running Pfam locally. The overlap with Pfam domain annotations were calculated by taking into account at least one overlapping position between motif and domain regions.

#### Random Forest classifier

From the python scikit-learn library [68], RandomForestClassifier class was applied to train and build a model. Entropy was selected as criterion, maximum tree depth was set to 4, number of features considered when looking for the best split was 4, and the number of trees in the forest was 10000. Balanced weight was applied for the training dataset by using the inverse proportion of class frequencies. For model training, the negative set was established by randomly selected 10000 sequences of equal residue length with the known instances, which is 8, from the human proteome. To get the feature importances from the trained classifier the feature_importances_ function (with default parameters) of the scikit-learn python package was used. For ROC curve analysis Stratified K-Folds cross-validator was carried out by using the scikit-learn StratifiedKFold library, which returns stratified folds preserving the percentage of samples for each class. This resulted in four folds of 18 and one fold of 19 positive samples (91 in total), and 2000 random samples for each five fold. The ROC curve calculation was based on the calculated confidence scores of hits (see below: Confidence score of predicted instances). The Depiction of the ROC curve analysis was created by the RocCurveDisplay function of the scikit-learn package.

#### Generating filtered human proteome for prediction

The human proteome set consisted of 20504 proteins for which Alphafold2 models were available. From this dataset, proteins with Secreted UniProt annotation (term was, subcellular location: Secreted) were filtered out, resulting in 18391 kept proteins. In the filtered human proteome, regions with Transmembrane/Extracellular UniProt region description were ignored during the prediction.

#### Confidence score of predicted instances

The trained classifier was applied to compute the confidence score of the 896 motif instance. Each predicted instance was scored by how many of the 10,000 random decision trees of the forest classify the given putative motif as binder. The ROC curve analysis was based on the calculated confidence scores.

### Integrating high throughput data and GO annotations

We collected the ciliary-centrosome system proteins from nine high-throughput (HT) proteomic studies [32–40]. Based on localization data of the experimental results, all proteins that were assigned a centrosomal or ciliary annotation in at least one HT study were collected. In total, 2213 and 958 proteins were collected for the ciliary and centrosomal systems, respectively. GO annotation based ciliary-centrosome system proteins were collected from the PANTHER classification system [27]. Proteins with Centrosome and/or Cilium GO Cellular component terms were considered as Ciliary-centrosomal proteins and were used in the analysis. Similarly, dynein system proteins were downloaded using the Dynein complex GO Cellular component term. In total, 634, 738 and 55 proteins were collected based on GO annotations for the centrosomal, ciliary and dynein systems, respectively.

### Enrichment

GO enrichment analysis was carried out by the GOC webserver (http://geneontology.org/). Overrepresentation test (Fisher’s exact test) of GO terms (all three aspects: biological process, molecular function, cellular component) was applied for the 674 LC8 binding partners, which included both the known and predicted proteins (four proteins, ATOSA BLTP3B, ENTREP1, ERCC6L2-AS1 were unmapped and excluded from the analysis by the GOC server). The test was performed with the integrated PANTHER Classification System [27] (current release was 17.0). False discovery rate (FDR) calculation based correction was applied. Enriched terms with FDR p-values above 0.01 were filtered out. Also, non-specific, generic terms (more than 1000 annotations) have been discarded.

### Evolutionary analysis

To compile evolutionary data for each protein containing LC8 binding motif, we created a dataset of orthologous sequences. These sequences were obtained by running the GOPHER prediction algorithm against the QFO database with default settings [69,70]. Subsequently, we performed multiple sequence alignments of the orthologs for each protein using the MAFFT algorithm with default parameters [71]. To classify the ortholog sequences, we utilized the UniProt taxonomic lineage, employing the five main evolutionary levels, Mammalia, Vertebrata, Metazoa, Fungi, and Eukarya, to determine the most specific term for each sequence. Then, protein level conservation of each LC8 partner was defined based on the orthologs with the most distantly related taxonomic term. For the evolutionary analysis, a minimum of three predicted orthologs was necessary at each level.

Based on the multiple sequence alignments of orthologs, we analyzed each aligned instance of the predicted LC8 binding motifs in terms of PSSM-based conservation. The PSSM score for each orthologous motif was calculated and then normalized using the human motif PSSM score as a reference. Subsequently, for each taxonomic level, we computed an average conservation score based on the normalized PSSM scores of the orthologous motifs belonging to that level. The calculated conservation scores of evolutionary levels were then used to determine motif conservation. An LC8 binding motif was considered conserved at a given level if the evolutionary level score exceeded 0.25. However, for four ciliary-centrosomal proteins (CCDC61, CFAP251, CATSPERZ, ZBBX), their LC8 binding motifs did not show conservation even in mammals. Due to this very weak conservation, these proteins were excluded from the analysis as it suggests that these motifs are likely false positives.

### Experimental procedure

#### Plasmid constructs

Truncated (1-270) coding sequences of OFD1, were obtained by PCR from cDNA library isolated from HEK293 cell line. The OFD1_1-270_ sequence was inserted into the in-lab modified bacterial expression vector pET-MBP (containing an N-terminal MBP-tag and a C-terminal 6HIS tag). The sOFD1-ΔTQT mutant was generated by substituting the core TQT triplet of the LC8 binding motif to alanins using quick-change mutagenesis. pEV vector including LC8 (DYNLL2) with an N-terminal 6xHis-tag was used for LC8 expression. For the pull-down experiments, full length DYNLL2 was cloned to pETARA vector (modified pET vector, with N-terminal GST-tag).

#### Protein expression and purification

6xHis-tagged LC8, GST-LC8 and N-terminal MBP, C-terminal 6xHis-tag-fused sOFD1 variants were expressed in E. coli BL21 (DE3) Rosetta cells (Novagene). Cultures were grown in LB-media, supplemented with 100 μg/ml ampicillin, 30 μg/ml chloramphenicol, at 37 °C until an optical density of 2.5 - 3 McFarland unit was reached. Overnight expression at 18 °C was induced by 0.1 - 0.3 mM IPTG. 6xHis-tagged LC8 was purified using Ni Profinity IMAC Resin (BioRad) followed by anion-exchange chromatography on HiTrap Q-HP column (GE Healthcare), and a final step of size-exclusion chromatography on Superdex 75 10/300 column (GE Healthcare).

#### GST pull-down experiments

For the pull-down experiments, GST-fused LC8 was used as bait, and MBP-, 6x-His-fused OFD1 variants as prey. For controls, GST and MBP proteins were used. The proteins were expressed in 100 ml cultures of E. coli BL21 (DE3) Rosetta cells as described above. Cells were harvested, resuspended in 5 ml PBS supplemented with 1 mg/ml lysozyme and 1 mM DTT, sonicated and clarified by centrifugation. 20 μl of Glutathione Sepharose 4B (Macherey-Nagel) (75%) resin was used for each experiment. Baits were incubated with the resin for 1 hour at room temperature, then washed with PBS supplemented with 1 mM DTT. Prey proteins were added and incubated for one hour. The beads were washed and eluted by boiling in a 2x SDS sample buffer and analyzed in tricine SDS PAGE [72].

#### Cell culture conditions

The hTERT-RPE1 cells (ATCC CRL-4000) were cultured in DMEM medium (Lonza) supplemented with 10% fetal bovine serum (Lonza) and Streptomycin-Penicillin (Lonza) in a 5% CO_2_ atmosphere at 37 °C.

#### Immunofluorescence

hTERT-RPE1 cells were seeded on poly-lysine-coated 21 mm coverslips. After 24 hours, cells were serum-starved for an additional 72 hours to generate ciliated cells. In case of non-ciliated cells, cells were fixed 48 hours after seeding. Cells were washed with DPBS without Ca^2+^ and Mg^2+^ (Lonza) two times and once in cytoskeletal buffer (10 mM PIPES, 100 mM NaCl, 3 mM MgCl_2_, 300 mM Sucrose, 5 mM EGTA, 0.5% Triton-X 100, pH 6.9) [73]. Subsequently, cells were fixed in a cytoskeletal buffer containing 4% PFA for 10 minutes at 37 °C. Fixed cells were washed with DPBS and permeabilized for an additional 30 minutes in 0.5 % Triton-X 100 in DPBS and incubated for 1 hour in a blocking buffer (2% BSA, 0.1% Triton-X 100 in DPBS) at room temperature. Primary and secondary antibodies were diluted in a blocking buffer. The cells were incubated with rabbit anti-OFD1 1:200 (Novus Biologicals NBP1-89355), goat anti-LC8 1:100 (Invitrogen PA5-47957), mouse anti-acetylated-α-tubulin 1:15000 (Sigma Aldrich 5335S) or anti-γ-tubulin 1:5000 (Sigma Aldrich T6557) overnight at 4 °C. Finally, cells were treated with secondary antibodies (anti-goat-Alexa Fluor 488 1:800, anti-mouse-CY3 1:800, anti-rabbit-Alexa Fluor 647 1:800 Jackson ImmunoResearch) for 1 hour at room temperature. Coverslips were mounted using ProLong Diamond antifade mountant with DAPI (Invitrogen) and imaged with an inverted Zeiss LSM 800 (Zeiss), the image processing was completed with Fiji ImageJ software [74]. The calculations of the colocalization were performed using the JACoP plugin of ImageJ software [75]. The Pearson’s coefficient and the Manders coefficient were calculated on Z-stack images of primary cilia and centrosomes of hTERT-RPE1 cells using manual thresholds. For the statistical analysis of the colocalization a randomization based Costes method was used.

#### Synthetic peptides

The synthetic peptides containing the 8-residue long LC8 binding motifs were ordered from Genscript Ltd. For better solubilization, the peptides containing the flanking regions of the motif extended their length to N and C terminally by 3–4 residues. The synthetic peptides were dissolved in HEPES or PBS buffer at 5 mM concentration, depending on the further used assay.

#### Fluorescence polarization

Fluorescence polarization experiments were carried out at 25 °C in PBS supplemented by 2 mM TCEP and 0.05% Brij using a Synergy H4 (BioTek Instruments) plate reader. N-terminally fluorescein 5-isothiocyanate (FITC) labeled, known binding peptide of LC8, BMF (FITC-TSQEDKATQTL) was used as a reporter peptide to form LC8-BMF complex. Briefly, 50 nM labeled BMF peptide was mixed with LC8 in a concentration to achieve 80% saturation, determined by direct titration (12 μM at 80% saturation). Subsequently, the above-mentioned complex was titrated with a two-fold dilution series of competitor ALMS1, AKAP9, OFD1, and CDK5RAP2 peptide and the decrease of polarization was recorded. To determine the K_d_ of the competitor peptide-LC8 complex, first the K_d_ of the BMF-LC8 complex was calculated. The polarization signal recorded at direct titration of BMF peptide with LC8 was plotted against the concentration of LC8. Then the K_d_ of the BMF-LC8 complex was calculated by the fitting of a direct binding equation in Origin 8. To determinate the K_d_ of the competitor peptide, the measured polarization was plotted against the concentration of competitor peptides in the competitive titration and the data was fitted by the competitive binding equation in Origin 8 where K_d_ of the BMF-LC8 complex was one of the parameters of the competitive equation. The titrations were carried out in triplicates in a 384 well plate format.

#### Surface Plasmon Resonance

Surface Plasmon Resonance experiments were carried out at 25 °C in a HEPES buffer (50 mM HEPES, 100 mM NaCl, 2 mM TCEP, 0.05% Tween-20 at pH 7.4) using a ProteOn XPR36 SPR instrument (Bio-Rad) on an HTG Ni-NTA sensor chip (Bio-Rad). The sensor chip was initialized according to the protocol of the manufacturer and activated using 100 mM NiCl_2_ solution. After the activation, 6xHis tagged LC8 was immobilized in 3 steps on the L1-L3 channels at 0.5 μM, 0.25 μM, and 0.125 μM concentration respectively until 1500 RU has been reached on the L1 channel. The L4 channel was used for reference channel without immobilized 6xHis tagged LC8. The motif containing peptides (OFD1, IQUB, CFAP91, MAP4, CEP152) were diluted in HEPES buffer and injected to the A1-A5 channels in two-fold dilution series starting at 30-50 μM concentration. At the A6 channel, HEPES buffer was injected as negative control. The final sensorgrams were generated by double referencing as the subtraction of the blank injection and the reference channel. The on rate (k_a_) and off rate (k_d_) constants were determined by using a 1:1 Langmuir equation fitted on the sensorgrams, the K_d_-s for the individual dilution series of the various immobilized LC8 surfaces were calculated as the quotient of the on and off rate constants.The overall K_d_ of the interactions was determined as the mean of the individual calculated K_d_ values of the three dilution series of peptides on the different LC8 surfaces.

## Acknowledgments

We thank Dr Gergő Gógl and Dr Ágnes Tantos for their valuable comments on the manuscript. The pEV vector including LC8 (DYNLL2) and the pETARA vector were the generous gift of Dr Péter Rapali and Dr László Nyitray. We thank Dr András Málnási-Csizmadia for the hTERT-RPE1 cell line.

## Authors contribution

Conceptualization, M.P., T.S. and Z.D.; Data curation, T.S., M.P.; Formal analysis, M.P., T.S.; Funding acquisition, Z.D. and T.S.; Investigation, T.S., M.P. M.F. and Z.D.; Methodology, M.P., T.S.; Supervision, Z.D.; Visualization, M.P., T.S.; Writing—original draft, M.P., T.S. and Z.D.

## Data and materials availability

All data are available in the main text or the Supplementary Materials.

