## Supplementary Materials for "Entering new circles: Expansion of the LC8/DYNLL1 interactome in the ciliary-centrosomal network through system-driven motif evolution"

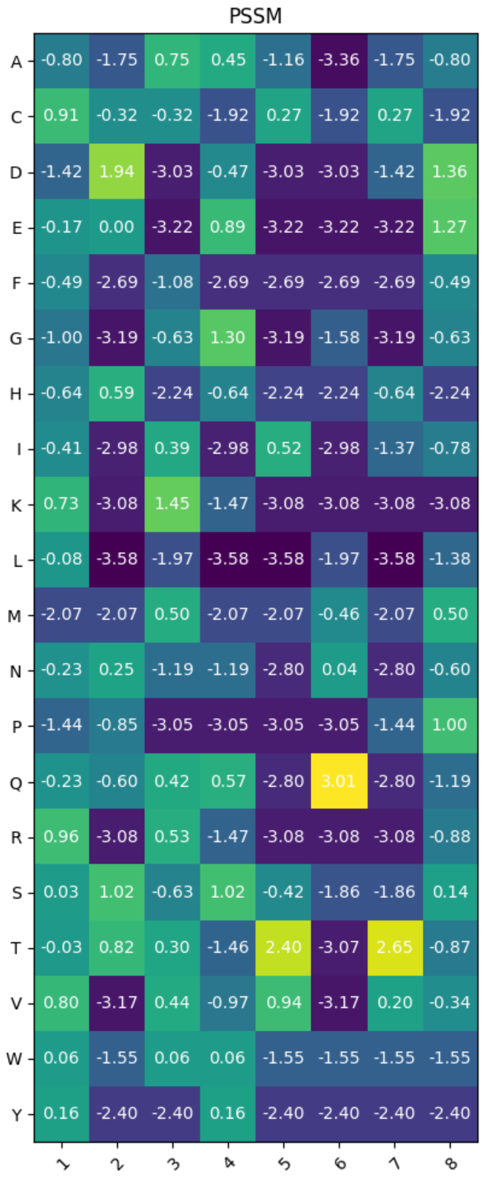


**Figure S1. Position Specific Scoring Matrix generated from the 91 known instances.**

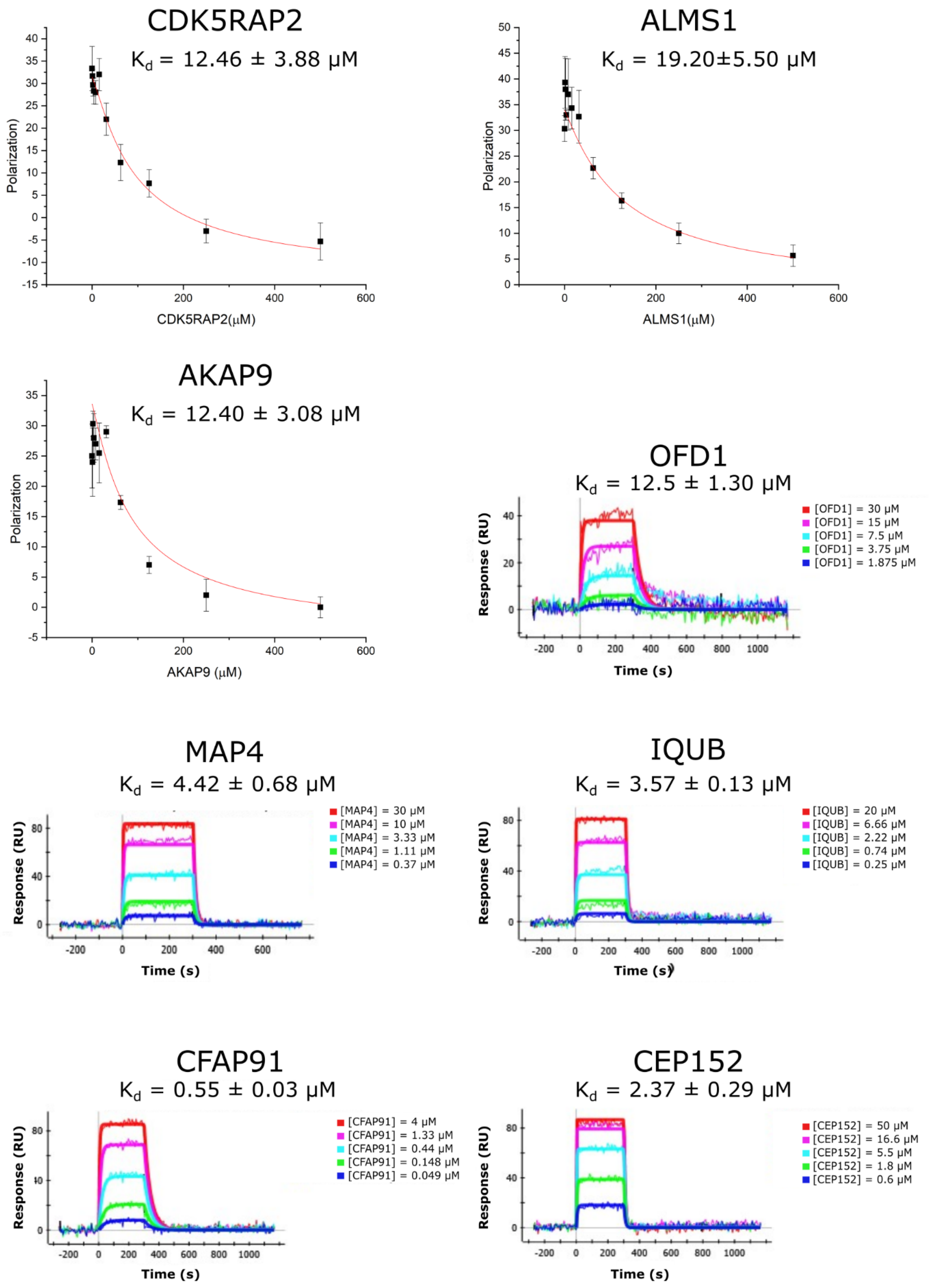


**Figure S2. Validation of LC8 binding partners on the motif level by fluorescence polarization (FP) and surface plasmon resonance (SPR).** Measurements were performed in triplicates (n = 3).

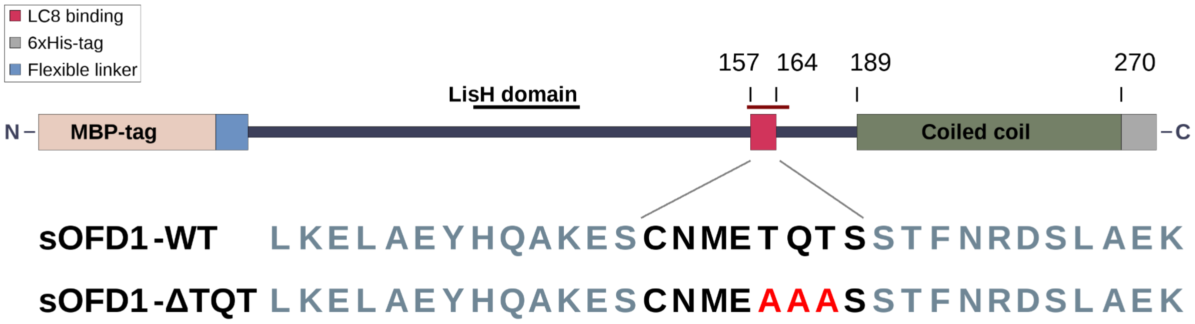


**Figure S3. Structure of the sOFD1-WT and sOFD1-ΔTQT constructs.**
